# A transposon surveillance mechanism that safeguards plant male fertility during stress

**DOI:** 10.1101/2020.10.03.324764

**Authors:** Yang-Seok Lee, Robert Maple, Julius Dürr, Alexander Dawson, Saleh Tamim, Charo del Genio, Ranjith Papareddy, Anding Luo, Jonathan C. Lamb, Anne W. Sylvester, James A. Birchler, Blake C. Meyers, Michael D. Nodine, Jacques Rouster, Jose Gutierrez-Marcos

## Abstract

Although plants are able to withstand a range of environmental conditions, spikes in ambient temperature can impact plant fertility causing reductions in seed yield and significant economic losses^1,2^. Therefore, understanding the precise molecular mechanisms that underpin plant fertility under environmental constraints is critical to safeguard future food production^3^. Here, we identified two Argonaute-like proteins whose activities are required to sustain male fertility in maize plants under high temperatures. We found that MALE-ASSOCIATED ARGONAUTE 1 and 2 (MAGO1 and MAGO2) associate with temperature-induced phased secondary small RNAs in pre-meiotic anthers and are essential to control the activity of retrotransposons in male meiocyte initials. Biochemical and structural analyses revealed how MAGO2 activity and its interaction with retrotransposon RNA targets are modulated through the dynamic phosphorylation of a set of highly conserved surface-located serine residues. Our results demonstrate that an Argonaute-dependent RNA-guided surveillance mechanism is critical in plants to sustain male fertility under environmentally constrained conditions by controlling the mutagenic activity of transposons in male germ cells.

## Introduction

Crop yield loss driven by high temperatures has been widely reported in many species, posing a real threat to food security^2,4,5^. Plant male reproductive development is especially sensitive to spikes in temperature^6^, with direct consequences on fertility and seed productivity^3,7^, likely due to metabolic and physiological changes resulting from epigenetic perturbations triggered by heat stress^8^. In rice, the temperature-sensitive genic male sterile mutants, *pms3* and *p/tms12-1*, have been found to carry mutations in long non-coding RNA (lncRNA) precursors that normally produce phased secondary small interfering RNAs (phasiRNAs)^9–11^ mainly in developing anthers. The slicing of rice lncRNA phasiRNA (*PHAS*) precursors is directed by two 22-nt miRNAs (miR2118 and miR2275) ^12,13^, leading to the formation of double-stranded RNA by RNA-DEPENDENT RNA POLYMERASE 6 (RDR6)^14^, which is sequentially processed by Dicer-like (DCL) proteins DCL4 or DCL5 into 21- or 24-nt phasiRNAs, respectively^15,16^. Interestingly, two rice Argonaute-like (AGO) proteins, AGO 18 and MEIOSIS ARRESTED AT LEPTOTENE1 (MEL1) have been found to be associated with 21-nt phasiRNAs and are required for male fertility ^13,17,18^. Thus, while the production of phasiRNAs in rice anthers has been linked to male fertility, the precise biological significance of this pathway in plants remains largely unknown.

It has been reported that maize anthers also accumulate two distinct populations of phasiRNAs, which occur at distinct phases of male germ cell development^19^. Of particular interest is the 21-nt phasiRNA subclass, which is abundant in the epidermis of pre-meiotic wild type anthers but absent in male-sterile mutants lacking a functional anther epidermis^19,20^. We postulated that the pre-meiotic phasiRNA pathway might play a critical role in supporting male fertility in plants. To this end, we scanned maize transcriptome data to identify components of the sRNA pathway specifically expressed in the epidermis of pre-meiotic anthers. We identified two Argonaute-like genes, with similarity to Arabidopsis AGO5 and rice MEL1, which we named Male-Associated Argonaute 1 and 2 (MAGO1 and MAGO2) (Fig S1A-B). Immunolocalization using specific antibodies revealed that MAGO1/2 accumulate primarily in the cytoplasm of epidermal cells of pre-meiotic anthers and in the nuclei of developing meiocytes (Fig. S1C).

To determine the possible roles of MAGO1/2, we first identified transposon insertion mutant alleles for both genes and found that while all wild-type and single mutant plants grown under field conditions produced fully viable mature pollen, double mutant plants were male-sterile (Fig. S2A-B). In addition, we generated *MAGO1/2* RNAi lines to simultaneously downregulate both genes (MAGO^KD^) (Fig. S2C) and found that most MAGO^KD^ lines displayed a range of male sterility phenotypes under field-grown conditions (Fig. 1A and Fig. S2D). Taken together, our genetic analyses revealed that both MAGO genes are required for male fertility in maize.

**Fig. 1.**
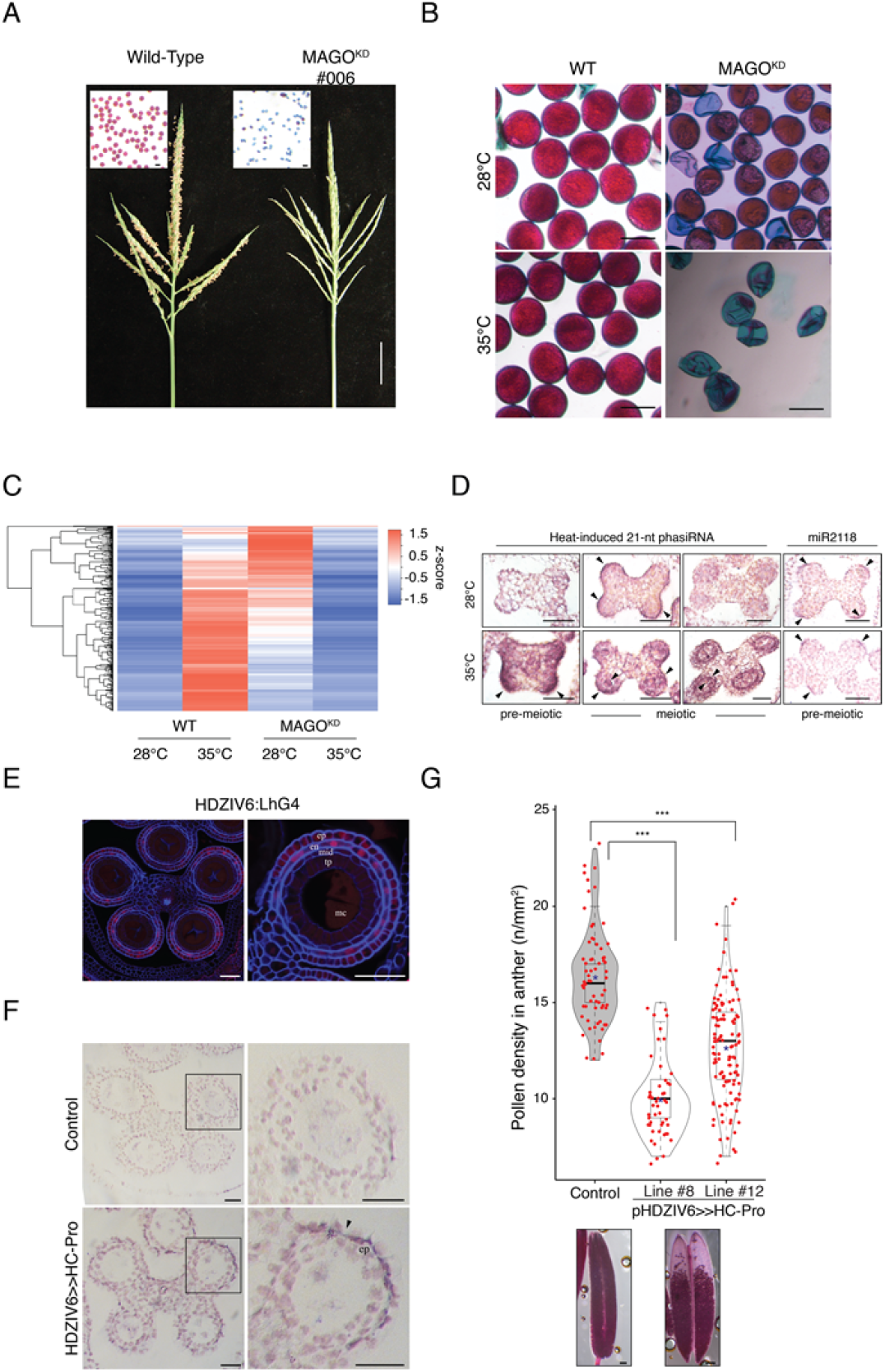
MAGO and pre-meiotic small RNAs are essential for male fertility in maize. (A) Male fertility defects observed in MAGO^KD^ plants grown under field conditions. Pollen grains after Alexander’s Staining and mature male inflorescences. Scale bars are 200 μm and 5 cm. (B) Pollen viability in wild-type and MAGO^KD^ plants under permissive conditions (28°C) and subjected to heat stress (35°C) before meiosis (n= 10). Scale bars are 200 μm. (C) Accumulation of 21-nt phasiRNAs in anthers of wild-type and MAGO^KD^ plants exposed to heat stress before meiosis. n ≥ 2 biologically independent samples each. (D) *in situ* RNA localization of 21-nt Hphasi and miR2118 in anthers of wild-type plants exposed to heat stress before meiosis. Scale bars are 50 μm. Black arrowhead indicates accumulation of sRNAs (E) Confocal images of NLS-dTomato reporter expression in anthers of HDZIV>>HC-Pro plants. Scale bars are 50 μm. ep, epidermis; en, endothecium; mid, mid-layer; tp, tapetum; mc, meiocyte. (F) Immunodetection of HC-Pro in anther epidermis in control and HDZIV>>HC-Pro plants. Scale bars are 50 μm. Black arrowhead, accumulation of HC-Pro. (G) Pollen viability in anthers from two independent HDZIV6>>HC-Pro lines (n≥30). Differences between groups were determined by one-way ANOVA, ***p < 0.001. Black lane, median; Red star, mean. Below, representative anthers after Alexander’s Staining. Scale bars 100 μm.

Because the spatial and temporal accumulation of MAGO1/2 mirrors the reported expression of 21-nt phasiRNAs and the predicted miR2118 trigger^19^, we next tested whether MAGO1/2 are involved in either the biogenesis or function of these phasiRNAs. We therefore performed immunoprecipitations using specific antisera and sequencing the bound small RNAs. We found that both MAGO1/2 are associated with the previously identified pre-meiotic 21-nt phasiRNAs (Fig. S3A-C), however, the abundance of these sRNAs was not significantly altered in MAGO^KD^ plants (Fig. S3D). This analysis revealed that MAGO1/2 are not directly implicated in the biogenesis of these particular phasiRNAs.

Given that male germline pre-meiotic phasiRNAs in rice are implicated in heat stress sensitivity and male fertility, we investigated the effects of heat stress on MAGO1/2 activity in maize. To this end, we grew wild-type and MAGO^KD^ plants under control temperature conditions and exposed them to a brief heat stress treatment either before or after male meiosis. Pollen viability was not affected in MAGO^KD^ plants when grown under non-stress (control) conditions, nor was it significantly affected in wild-type or MAGO^KD^ plants exposed to heat stress after meiosis (Fig. S4A). However, heat stress applied before meiosis had profound effects on pollen viability in MAGO^KD^ plants, becoming largely infertile, while wild-type plants remained relatively unaffected (Fig. 1B and Fig. S4B). Taken together, our genetic analyses revealed that both MAGO genes are required for male fertility under heat stress. Further, we postulated that a new class of MAGO-associated pre-meiotic phasiRNAs might determine male fertility under restrictive temperature conditions. To test this idea, we sequenced sRNAs from pre-meiotic anthers after a short exposure to heat stress (72 h/35°C). We found a massive accumulation of a discrete group of 21-nt phasiRNAs (Hphasi) in wild-type plants, but not in MAGO^KD^ plants (Fig. 1C and Fig. S5). Analysis of uncapped RNA ends provided strong evidence for miR2118-directed cleavage of Hphasi lncRNA precursors in pre-meiotic wild-type anthers (Fig. S6). The spatial distribution of these sRNAs was determined by RNA *in situ* localisation on anther sections from wild-type plants grown under permissive and restrictive conditions (Fig. 1D). We found that heat stress triggered Hphasi accumulation, primarily in the epidermis of pre-meiotic anthers and also in the inner anther cell layers at the onset of meiosis. However, we did not observe any increase in the accumulation of miR2118 in pre-meiotic anthers after heat stress. We therefore sought to determine whether the biogenesis of these RNAs play a role in male fertility. Given that HC-Pro can directly bind and sequester small RNAs^21,22^, we reasoned that HC-Pro induction in heat-stressed pre-meiotic anthers could interfere with the biogenesis and/or function of this particular class of phasiRNAs. Therefore to test this hypothesis, we first generated transgenic plants carrying a dexamethasone (DEX)-inducible viral gene-silencing suppressor – Helper Component-Proteinase (HC-Pro)^23,24^. We next immunoprecipitated HC-Pro from DEX-treated anthers and sequenced the bound sRNAs. We found that HC-Pro expression in pre-meiotic anthers was associated with 21-nt phasiRNAs and their predicted miR2188 trigger (Fig. S7A-D). To further define the role(s) of these sRNAs, HC-Pro was ectopically expressed in anthers before and after meiosis. Pre-meiotic induction resulted in a near complete lack of pollen grains at anther maturity, whereas post-meiotic HC-Pro induction exerted only minor effects on pollen production (Fig. S7E). To establish whether the HC-Pro-mediated effects on male fertility were specifically linked to sRNAs produced in anther epidermal tissues, we generated maize plants with HC-Pro under the control of an epidermal-specific HDZIV6 trans-activation system (HDZIV6>>HC-Pro) and confirmed exclusive HC-Pro expression in the anther epidermis by immunolocalization using a specific HC-Pro antibody that we generated (Fig. 1E-F). Notably, we found that the production of pollen grains in HDZIV6>>HC-Pro plants was drastically reduced compared to control plants (Fig. 1G). Collectively, these data provide evidence for the activation of a specific sRNA cascade in anther epidermis that is critical to support male fertility under restrictive temperature-stress conditions.

Computational genome-wide scans showed that Hphasi mapped primarily to transposable elements (TEs) and specifically to long terminal repeat retrotransposons (LTRs) within the maize genome (Fig. S8A-B). Because TEs are typically deregulated in plants during periods of genome shock or abiotic stress^25^, we reasoned that the HphasiRNAs might be involved in the silencing of TEs in anthers following environmental stress exposure. To test this hypothesis, we performed RNA sequencing (RNA-seq) to compare global changes in gene expression in pre-meiotic anthers of MAGO^KD^ and wild-type plants. Under permissive conditions, we did not observe any significant changes in the anther-specific expression of coding genes between WT and MAGO^KD^ (Fig. S8C). By contrast, exposure to heat stress before meiosis resulted in global changes in anther gene expression in all plants tested independent of genotype (Fig. S8C-D and Table S1). Notably, we found that LTR retrotransposons were significantly deregulated in anthers and in meiocytes of MAGO^KD^ plants after heat stress (Fig. 2A-B). In addition, most of the retrotransposons upregulated in heat-stressed MAGO^KD^ plants were of the low-copy Gypsy (LRGs) class predicted to be targeted by Hphasi (Fig. S9A and Table S2). Subsequent analysis of uncapped RNA ends provided strong support for the directed cleavage of LRGs by Hphasi (Fig. 2C and Fig. S9B). To test if MAGO and associated phasiRNAs are involved in modulating the mobility of these LRGs, we carried out transposon display and retrotransposon-sequence capture on progenies generated from reciprocal crosses between wild-type and MAGO^KD^ plants grown under permissive conditions. These analyses revealed retrotransposons to be highly mobile only in the male germ cell lineage of MAGO^KD^ plants (Fig. 2D and Table S3).

**Fig. 2.**
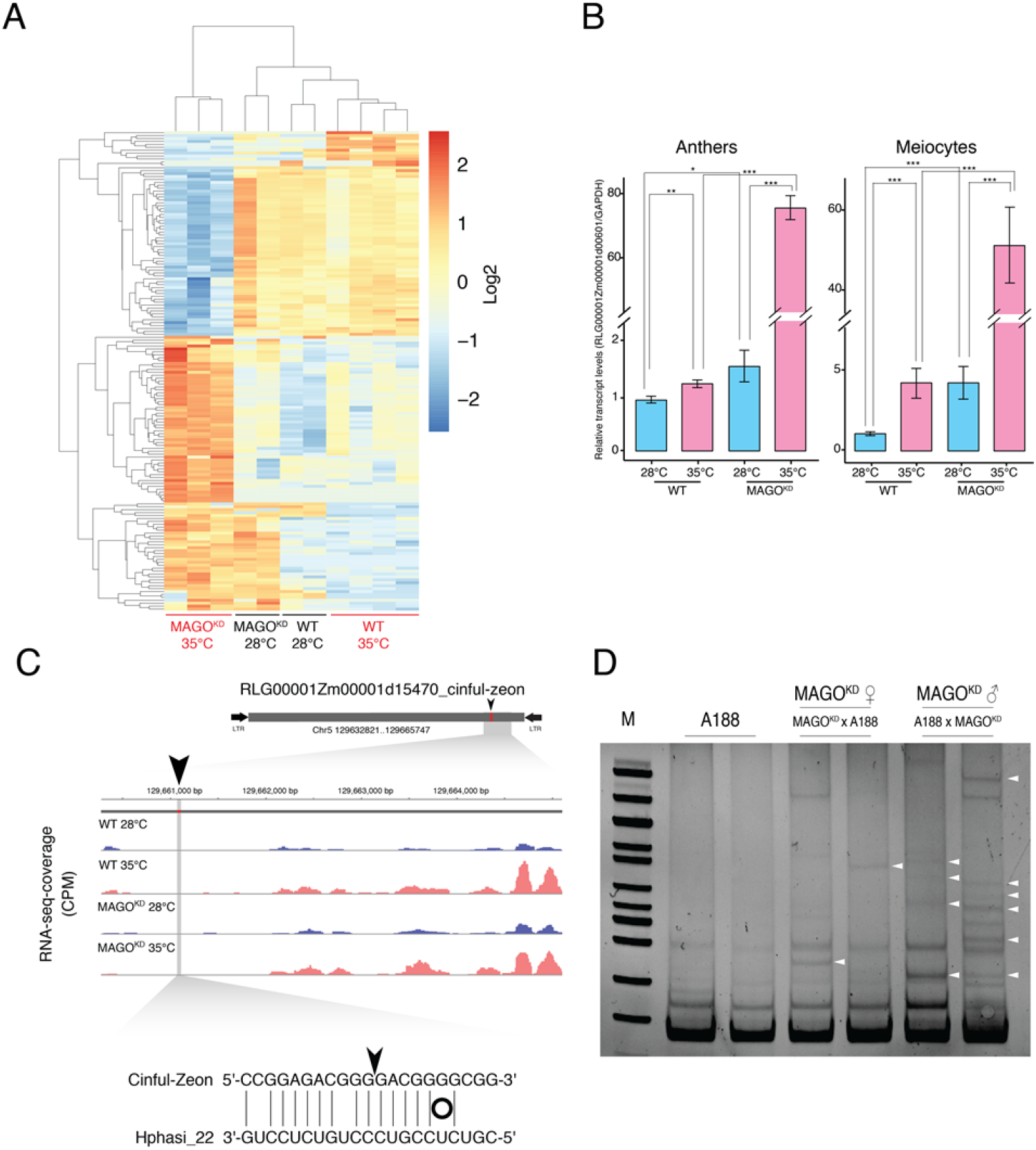
MAGO1/2 are necessary to silence stress-activated retrotransposons in maize male germ cells. (A) Heatmap analysis of transposon transcripts deregulated in anthers of wild-type and MAGO^KD^ plants exposed to heat stress before meiosis. n ≥ 2 biologically independent samples each. (B) Relative expression of a MAGO regulated LTR in anthers and meiocytes of wild-type and MAGO^KD^ plants exposed to heat stress before meiosis (n= 4). Differences between groups were determined by one-way ANOVA, *p < 0.05; **p < 0.01; ***p < 0.001. Black lane, median; Red star, mean. (C) Coverage of RNA-seq of retrotransposons targeted by Hphasi in anthers of wild-type and MAGO^KD^ plants exposed to heat stress before meiosis. n ≥ 2 biologically independent samples each. Black arrowhead and red box indicate the location of predicted sites for Hphasi-directed slicing remnants. n= 3 independent samples each. Grey box, retrotransposon; Black arrow, long terminal inverted repeats. (D) Transposon display showing the presence of retrotransposon insertions in progenies from wild-type (A188) plants and from reciprocal crosses with MAGO^KD^. Each lane represents a pool of 25 independent plants. n= 2 biologically independent experiments. White arrowheads indicate new retrotransposon insertions.

Our data thus far point to a role for MAGO proteins in protecting male germ cells from the deregulated activity of retrotransposons that occurs during heat stress. In order to understand how MAGO activity is regulated by heat stress, we first analysed our RNAseq data. However, we did not observe significant changes in transcript levels in response to heat stress, indicating that MAGO activity may be modulated post-translationally. Indeed, previous studies in maize and rice have identified other Argonaute-like proteins that are phosphorylated in anthers^26,27^. We therefore performed a phosphoproteome analysis of pre-meiotic wild-type anthers and identified major changes in phosphorylation induced by short exposure to heat stress (Fig. 3A and Table S4). Specifically, a region in the PIWI domain of MAGO2 that contains four serines (S989, S990, S994 and S998) and one threonine (T995) residue was significantly hypo-phosphorylated in response to heat stress (Figure S10). To further understand the biological significance of these dynamic changes in MAGO2 phosphorylation, we modelled the structure of the conserved catalytic domain of MAGO2 and analysed the electrostatic potential of the molecular surface of different phosphorylation variants (Fig. 3B and Movie). We found that the residues targeted by phosphorylation are located at the surface of a loop following the PIWI domain (Fig. S11A). This finding is in line with the conserved presence of phosphorylated serine/threonine residues in the PIWI loops of other catalytically active plant and animal Argonautes (Fig. S11B and Table S5). These phosphorylated residues are in close proximity to the central cleft and to an L2 loop, which are considered important for the regulation of sRNA:target interactions^28^. Notably, we found that the hyper-phosphorylation of S989-S998 does not cause significant changes to MAGO2 structure, however is predicted to cause profound changes to the electrostatic surface potential of the upper central cleft region (Fig. 3B). To test if these stress-induced changes could affect the affinity for sRNAs and/or the enzymatic activity of MAGO2, we immunoprecipitated MAGO2 from anthers of plants exposed to a pulse of heat stress prior to meiosis and quantified the bound small RNAs. While we did not find significant differences in the association of MAGO2 with 21-nt Hphasi under control and heat-stress conditions (Fig. 3C), the abundance of retrotransposon RNA targets associated with MAGO2 was significantly reduced after exposure to heat stress (Fig. 3D). Because of this, we reasoned that the changes in surface charge brought about by MAGO2 hypo-phosphorylation could regulate the interaction between sRNA and target to sustain the rapid silencing of deregulated retrotransposons in the male germ cells. To test this hypothesis, we generated an *in vivo* reporter system to test the catalytic activity of wild-type MAGO2 and a series of phosphoresistant (S>A) and phosphomimetic (S>E) MAGO2 mutants (Fig. S12). We also generated mutations in MAGO2 residues predicted to be essential for sRNA binding (Y676E) and small RNA-directed RNA cleavage (D835E) for inclusion in this experiment. We found that while sRNA binding was affected in MAGO^Y676E^, it was not significantly affected in either MAGO^S>A^, MAGO^S>E^ or MAGO^D835E^ (Fig. 3E). Silencing activity was, however, abolished in MAGO^Y676E^ and significantly reduced in only MAGO^S>E^ (Fig. 3F). These data suggest that stress-mediated hypo-phosphorylation of MAGO2 is important for its effective interaction with retrotransposon RNA targets. This model is consistent with previous studies in humans, which have shown that hyper-phosphorylation of the PIWI domain of AGO2 (S824–S834) impairs RNA:target association while hypo-phosphorylation expands the target repertoire ^29,30^.

**Fig. 3.**
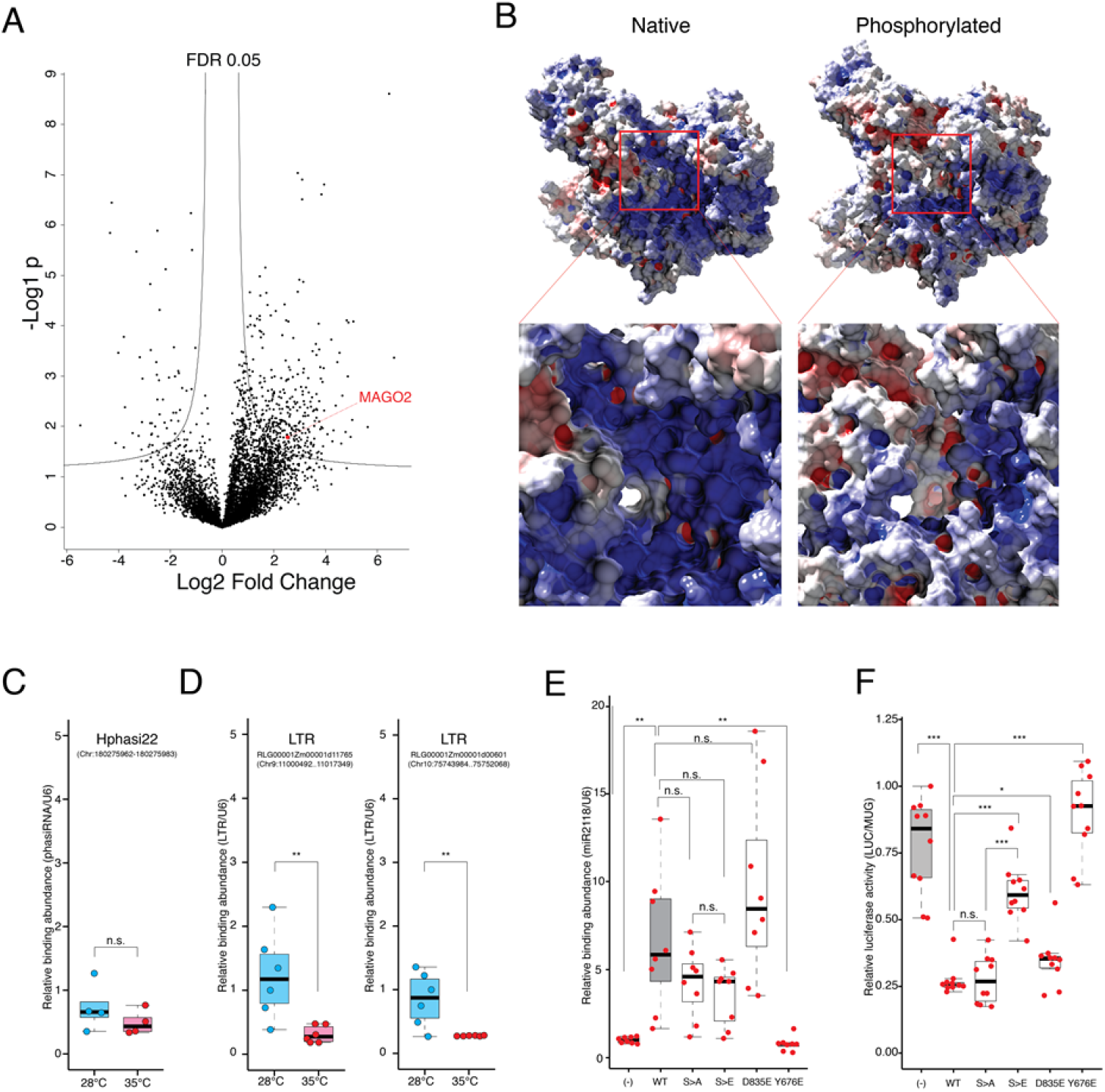
MAGO activity is modulated by dynamic changes in phosphorylation induced by heat stress. (A) Volcano plot showing dynamic changes in phosphorylation in premeiotic anthers after heat stress and identification of MAGO2 as a differentially phosphorylated protein. Threshold FDR<0.05; Log2 fold change>1.5; n= 6 independent biological replicates. Red circle, MAGO2 phosphopeptides. (B) Electrostatic potential distribution on the molecular surface of native and phosphorylated MAGO2 showing negative (red), positive (blue) and hydrophobic (grey) residues. Red boxes-close views of the central cleft and L2 loop known to be implicated in the regulation of sRNA:target interactions. (C) Accumulation of Hphasi22 bound to MAGO2, shown by immunoprecipitation coupled to reverse-transcriptase polymerase chain reaction (qRT-PCR), in pre-meiotic anthers from wild-type plants exposed to control (28°C) and heat-stress conditions (35°C). n = 4 independent biological replicates. Differences between groups were determined by paired t-test, n.s. not significant. (D) Accumulation of retrotransposon RNA bound to MAGO2, shown by immunoprecipitation coupled to reverse-transcriptase polymerase chain reaction (qRT-PCR), in pre-meiotic anthers from wild-type plants exposed to control (28°C) and heat-stress conditions (35°C). n= 6 independent biological replicates. Differences between groups were determined by paired t-test, **p < 0.01. (E) Silencing activity of MAGO2 in wild-type and MAGO2 mutants for residues implicated in differential phosphorylation (S>A and S>E), and in residues predicted to be necessary for target cleavage (D835E) and sRNA binding (Y676E). n= 10 independent biological replicates. Differences between groups were determined by one-way ANOVA, *p < 0.05; ***p < 0.001; n.s. not significant. (F) Abundance of bound sRNAs in wild-type MAGO2 and mutants, shown by immunoprecipitation coupled to reverse-transcriptase polymerase chain reaction (qRT-PCR). n= 10 independent biological replicates. Differences between groups were determined by one-way ANOVA, **p < 0.01; n.s. not significant.

In sum, we have identified in maize two Argonaute-like proteins acting alongside a distinct group of sRNAs in a male germline-specific manner. This pathway is remarkably similar to the small RNA-mediated recognition pathway found in mammalian male germ cells, which acts to silence retrotransposon activity^31^. Unlike in animals, transposon activity is however more permissive in plants. For instance, bursts of transposon mobility have helped shape plant genomes and alter transcriptional responses^32^. Further, the insertion of retrotransposons in gene regulatory regions have caused notable dramatic effects on plant phenotypes – a property that was repeatedly exploited during the process of crop domestication^33^. Nevertheless, plants must regulate retrotransposon activity because, as shown for genomic shock, deregulated retrotransposon activity often results in genomic and epigenomic instability, and ultimately causes negative effects on offspring fitness^25^. Our findings reveal an Argonaute-mediated pathway that protects plant male fertility by acting as a pre-meiotic surveillance mechanism activated in the somatic cells that surround the germ cell precursors. The biological significance of this molecular pathway operating under heat stress conditions to restrict retrotransposon activity in developing male germ cells may be to prevent the widespread transmission of unwanted or deleterious mutations in wind-pollinated plants such as grasses^34–36^. The manipulation of this molecular mechanism in crops will undoubtedly become a useful strategy in future efforts to enhance male fertility and sustain seed productivity under unpredictable and stressful climate conditions.

## Acknowledgements

We thank Gary Grant and Peter Watson for help with plant husbandry; Liliana M. Costa for discussions and comments on the manuscript. This research was supported by awards from US National Science Foundation (1027445 to A.W.S. and 1649424 & 1754097 to B.M.), European Research Council (Grant 637888 to M.D.N.) and BBSRC (BB/L003023/1, BB/N005279/1, BB/N00194X/1 and BB/P02601X/1) to J.G-M.

## Author Contributions

Y-S. L. and R.M. cultivated plants, harvested samples and collected phenotypic data. R.P., JG-M and J.R. identified transposon insertions. J.D. performed the phosphoproteome analysis. R.M. and Y-S. L. performed immunoprecipitation and small-RNA-seq libraries. A.L. and J.L. generated constructs and transgenic lines for HC-Pro and HDZIV6 transactivation. R.P. constructed nanoPARE libraries. R.M, A.D, S.T. and M.D.N. performed the bioinformatic analysis. C. de G. performed the molecular dynamics simulation and protein modelling. Y-S. L., R.M., Y.D, A.D., S.T., C. de G. prepared figures and tables. A.W.S, J.B., B.C.M., M.D.N., J.R. and J.G-M. co-ordinated experiments. Y-S. L. and R.M. and J.G-M. conceived the project. J.G-M. wrote the manuscript with input from the rest of the authors.

## Data and Materials Availability

Sequence data (mRNA-seq, nanoPARE-seq, sRNAseq and LTR-seq) that support the findings of this study have been deposited at the European Nucleotide Archive (ENA) under the accession code ERP118841.

## List of Supplemental Materials

Materials and Methods

Figures S1-S11

Tables S1-S7

Movie 1

