## Supplemental Data for "A transposon surveillance mechanism that safeguards plant male fertility during stress"

**Supplementary Materials**

### 1. Materials and Methods:

#### Plant material and growth conditions

The maize cultivar A188 was used in all experiments. Plants were grown at 28°C day/ 20°C night in a 16 hr light/8 hr dark cycle with a light intensity of 230 µE m-2 s-1. Seeds were germinated in three-inch diameter pots containing peat-based soil and grown for three weeks before transferring to eleven-inch or 40 L pots to flowering. Anthers were manually dissected at v12 stage and meiotic stage was determined by acetocarmine staining. For heat stress experiments, maize plants grown in eleven-inch pots were transferred to growth chambers for 3 days (16 h light at 35°C/ 8 h dark at 25°C, light intensity 230 µE m-2 s-1, humidity 75%). Tobacco plants (*Nicotiana benthamiana*) were grown on M2 soil (Levington Advance, UK) at 22°C in a 16 h light/ 8 h dark cycle with light intensity of 100 µE m-2 s-1.

#### Identification of transposon insertion mutants

To identify transposon insertion lines for Zm00001d007786 (GRMZM2G05903) and Zm00001d013063 (GRMZM2G123063) we screened a Mutator insertion mutant population generated by Biogemma, an Ac/Ds mutant population (*1*) and an UniformMu mutant population (*2*). Insertion lines were confirmed by PCR (Supplementary Table S6) and backcrossed to A188 inbred for four generations before analysis.

#### Vector construction and generation of transgenic plants

We generated a MAGO1/2-RNAi vector by subcloning a portion corresponding to position 2151-2400 of Zm00001d007786 (GRMZM2G05903) and a fragment corresponding to position 1001-1250 of Zm00001d013063 (GRMZM2G123063) into a pCsVMV::intOsActin-intStLS1-terSbHSP vector using Golden Gate cloning.

To generate a chemically inducible helper component-proteinase (HC-Pro) construct we synthetized a 2,211 bp fragment from the Wheat Streak Mosaic Virus WSMV containing Gateway Recombination flanking sequences (Invitrogen, USA). The synthetic fragment was subcloned into a two-component dexamethasone-inducible expression binary vector named pZZ-TOP, which is derived from pTF101.1 and carries a LhG4:GR synthetic gene fusion and a bi-directional pOp6 promoter (*3*). This vector enables the co-expression of a Beta-Glucuronidase (GUS) reporter and HC-Pro after exposure to 20mM Dexamethasone (DEX).

To enable the epidermal expression of HC-Pro, we generated a pZmHDZIV6-LhG4 vector using MultiSite Gateway Recombination (Invitrogen, USA). The native 3164 bp promoter and the 945 bp native terminator of Zm00001d002234 (GRMZM2G001289) was cloned into pDONR221 P1-P4 and pDONR221 P3-P2, respectively. The maize codon-optimized LhG4 was cloned into pDONR221 P4r-P3r. The resulting three entry vectors were then recombined into the binary vector pAL010, a derivative of pTF101.1 carrying a Gateway Recombination Cassette and a bi-directional pOp6 promoter enabling the co-expression of a Beta-Glucuronidase (GUS) and NLS-tdTomato reporters. All constructs were fully sequenced before transformation in maize using *Agrobacterium tumefaciens* strains LBA4404 (MAGO1/2-RNAi and pZmHDZIV6-LhG4) or EHA101 (HC-Pro).

To determine the in vivo activity of MAGO proteins we generated a firefly Luciferase silencing reporter system. First, we generated a construct containing a nopaline synthase (NOS) promoter (pNOS), a firefly luciferase (FLUC) fused to four miR2118-target sequences (PHAS), a truncated-GFP (ΔGFP) and a NOS terminator (tNOS). This synthetic fragment was cloned in pBINPLUS using *Hin*dIII and *Bam*HI restriction enzyme digestion. The miR2118 target region was designed to have optimal hybridization energies with the RNA target. The resulting construct was digested with *Sma*I and *Eco*RI to enable the insertion of a p35S::GUS:tNOS fragment derived from the pSLJ4J8 vector. We also generated a fragment containing the octopine synthase (OCS) promoter, an artificial miRNA based on maize miR2118c (amiR2118) and a nopaline synthase (NOS) terminator. This fragment was cloned into pBINPLUS using *Asc*I restriction enzyme digestion. To simultaneously express MAGO and amiR2118, we generated codon-optimized MAGO1 and 2, containing a FLAG tag in the carboxy-terminal end, which were cloned into pCsVMV::intOsActin-terSbHSP using SapI resulting in the binary vectors pBIOS11743 and pBIOS11746, respectively. For the phosphorylation study, we generated codon-optimized MAGO2 phosphomimetic (S>E) and phosphoresistant (S>A) forms, containing a FLAG tag at the carboxy end, which were cloned into pCsVMV::intOsActin-terSbHSP using SapI resulting in the binary vectors pBIOS11747 and pBIOS11748, respectively. MAGO2 mutants for sRNA binding (Y676E) and cleavage (D835E) were generated using Q5® Site-Directed Mutagenesis Kit (NEB, UK). The pOCS::amiR2118:tNOS was sub-cloned into pBIOS11743 and pBIOS11746 using AscI restriction enzyme digestion. Generated constructs were fully sequenced and transformed in *Nicotiana benthamiana* using *Agrobacterium tumefaciens* GV3101.

To study the in vivo localization of MAGO proteins, codon-optimised coding regions for each gene were subcloned in pGWB441 (*4*) to generate c-terminal EYFP protein fusion after Gateway Recombination (Invitrogen, USA). All constructs generated in this study were fully sequenced and transformed in *Nicotiana benthamiana* using *Agrobacterium tumefaciens* GV3101.

**Collection of maize meiocytes**

The meiotic stage of anthers was determined using acetocarmine staining. Meiocytes were isolated by manual micromanipulation in RNase-free PBS and collected using microglass pipettes controlled by UMP3 UltraMicroPump (WPI). Collected meiocytes were directly frozen in liquid nitrogen and stored at -80°C.

**Antisera preparation and immunopurification**

Polyclonal antisera were raised in rabbit against synthetic peptides for MAGO1 (VETEHQQGKRSIYRI) or MAGO2 (CVAAREGPVEVRQLPK) (Eurogentec, Liege, BE). To generate HC-Pro antisera, a partial DNA fragment was chemically synthesized (Integrated DNA Technology, UK) and cloned in pET29a (Novagen, Merck, Darmstadt, Germany). A soluble fraction of HC-Pro was isolated and purified by metal affinity chromatography and polyclonal antiserum was raised in rabbit (Eurogentec, Liege, BE). All antisera were affinity purified using a Sulpholink coupling gel system (Pierce, Rockford, IL).

**MAGO immunoprecipitation**

For immunoprecipitation, maize anthers were isolated by micromanipulation and ground in extraction buffer (20 mM Tris-Cl pH 7.5, 300 mM NaCl, 5 mM MgCl2, 5 mM DTT, 1% (v/v) cOmplete EDTA-free protease inhibitor cocktail (Merck, Darmstadt, Germany). The lysates were pre-cleared with protein-A agarose beads (Sigma-Aldrich, Poole, UK) and incubated with anti-MAGO1 (1:200), anti-MAGO2 (1:200) or anti-HC-Pro (1:100) antibodies for 2h at 4°C. Protein-A agarose beads were added to the sample and incubated for a further 2h. Subsequently, the beads were washed 4-5 times using extraction buffer supplemented with 0.5% NP-40 (Sigma,

St. Louis, MO). For western blot analysis, beads were suspended in 1x SDS loading buffer and heated at 95°C. For isolation of small RNA, the bead slurry was digested with proteinase K (100 μg·mL-1) and incubated for 1 h at 37°C prior to RNA extraction using TRIZOL® Reagent (Invitrogen, UK).

**RNA extraction**

Total RNAs were extracted from immature tassels, anthers, meiocytes or leaves with TRIzol® Reagent (Invitrogen, UK) as per manufacturer's instruction. RNAs for sequencing were extracted using Direct-zol RNA miniprep kit (ZYMO Research, Cambridge). The extracted RNAs were quantified using Nanodrop (ThermoFisher, UK) and the quality checked with a Bioanalyzer 2100 (Agilent, UK).

**Preparation of RNA libraries and sequencing analysis**

To isolate small RNAs (sRNAs), total RNA was fractionated on a 15% polyacrylamide TBE-Urea gel (Novex, UK). The gel was stained with ethidium bromide for 5 mins at room temperature and visualised with a UV illuminator. Gel pieces were macerated by incubation with 300 mM NaCl overnight followed by RNA precipitation. For sRNA sequencing, fractionated sRNAs were used to construct libraries using the TruSeq Small RNA Sample Preparation Kit (Illumina, UK) and sequenced in single-end 50 base mode an Illumina HiSeq platform (University of Delaware). For total RNA sequencing, total RNAs with RNA integrity number (RIN) >8.0 were used for library construction using a TruSeq RNA Sample Preparation Kit (Illumina, UK) and sequenced in single-end 150 base mode an Illumina NexSeq platform (University of Warwick). NanoPARE libraries were prepared as described previously (Schon et al. 2018). Briefly, cDNA libraries were generated from 5 ng of total RNA using the original Smart-seq2 protocol (Picelli et al. 2013). Five nanograms of cDNA was tagmented using Nextera DNA Flex library preparation kit as described in the manufacture instructions. Tagmented cDNA was purified using Zymo DNA Clean and Concentrator kit and eluted with 20ul nuclease-free water. This purified tagmented product was split into halves and used as a substrate for final enrichment PCR with either Tn5.1/TSO or Tn5.2/TSO oligonucleotide primer sets (Schon et al. 2018). PCR reaction products from Tn5.1/TSO and Tn5.2/TSO oligonucleotide primer sets were pooled together and purified using Beckman Coulter AMPureXP DNA beads. The nanoPARE libraries were sequenced in single-end 50 base mode on an Illumina Hi-Seq 2500 instrument. Library details and number of reads are provided in Table S7.

**Induction of HC-Pro in developing anthers**

Maize HC-Pro transgenic plants were grown to adult stage and whorls were cut open to reveal the developing male inflorescence (tassels). Each floret was filled with a DEX solution (50 µM DEX, 0.1% Silwet-77) and allowed to grow to maturity under normal growth conditions. To determine the efficiency of DEX-induction, anthers were submerged in GUS solution (50 mM sodium phosphate, 1 mM potassium ferricyanide, 1 mM potassium ferrocyanide, 0.1% Triton X-100, 10 mM EDTA, 1.0 mg·mL-1 5-Bromo-4-chloro-3-indolyl-β-d-glucuronic acid), incubated at 37°C for 16-24 h until blue precipitate was observed. DEX--treated and mock-treated anthers were sampled to determine pollen viability. To check induction of HC-Pro, total protein was extracted from anthers using QB buffer (100 mM KPO4 (pH 7.8), 1 mM EDTA, 1% Triton X-100, 10% glycerol, 1 mM DTT, Protease inhibitor cocktail (Roche, UK). The extracted proteins were quantified by Quick Strat Bradford Protein Assay (Bio-Rad, UK). Western blot detection was carried out with immunopurified anti-HC-Pro antisera and images recorded using an ImageQuant gel documentation instrument (GE Healthcare Life Sciences, UK).

**Confocal microscopy analysis**

To determine transcriptional activation of pZmHDZIV6::LhG4 in anther epidermis, transgenic plants were grown to reproductive stage and tassels were fixed in SR2200 solution (4% PFA in PBS (pH 7.4), 0.1% SR2000 (Renaissance Chemicals)), vacuum infiltrated, washed with PBS and submerged in ClearSee solution (10% xylitol (w/v), 15% sodium deoxycholate (w/v), 25% urea (w/v) (Kurihara et al. 2015). The samples were vacuum infiltrated, incubated until tassels were cleared, washed and stored in PBS. The cleared tissue was embedded in 4% low melting agarose in PBS and the embedded tissue was mounted onto vibratome blocks, 150 □m sections were cut by Lancer Vibratome Series 1000 (TPI, USA). Tissue slices were placed onto glass slides, covered with a coverslip and imaged with a LSM710 confocal microscope (Zeiss, Jena, GE).

**cDNA synthesis and RT-PCR**

Total RNA was treated with Ambion® TURBO DNase kit (Life technologies, USA). DNase-treated RNAs were used for cDNA synthesis using Superscript® reverse transcriptase II (Invitrogen, UK). Semi-quantitative RT-PCR was performed using templates as the synthesized cDNAs. GAPDH was used for data normalization. For Quantitative real-time RT-PCR (qRT-PCR) the optimal number of cycles was determined for each gene. PCR cycling conditions included denaturing at 95ºC for 15 s, annealing at 57ºC for 30 s and extension at 72ºC for 45, using a Bio-rad qRT-PCR machine (BioRad, UK). Changes in expression levels were calculated via the ΔΔCt method. To ensure primer specificity, qRT-PCR was done when the melting curve showed a single peak. To quantify small RNAs we used stem loop qRT-PCR following previously reported methods (Yang et al. 2014; Varkonyi-Gasic, 2017) with minor modifications. Reverse transcription was performed using the RevertAid First Strand cDNA Synthesis Kit (Thermo Scientific, USA) as per the manufacturer's instructions. For stem loop qRT-PCR, we used a 10 μL of RT reaction mixture containing 1 μL of RNA, 1 μL of RT primer (5 μM) and 1 μL of U6 RT primer (5 μM), 1 μL of 10 mM dNTP Mix, 2 μL of reaction buffer, 0.5 μL of Ribolock RNase inhibitor (20 U/μL), 0.5 μL revertAid M-MuLV Reverse Transcriptase (200 U/μL). The mixture was incubated at 25℃ for 5 min, and then incubation was continued at 42℃ for 60 min. The reaction was inactivated by heating at 70℃ for 5 min and to which 1 μL of RT product, 5 μL of SYBR Green real-time PCR Master Mix, and 1 μL of primer (forward and reverse, 1 μM each) was added. Reactions were incubated in a PCR cycler at 95℃ for 3 min, followed by 40 cycles of 95℃ for 5 s, 62℃ for 35 s. Primer sequences are listed in Supplementary Table S6.

**Display and high-throughput sequencing of retrotransposon insertions**

For the detection of retrotransposon insertions, we used Splinkerette PCR (*5*). Genomic DNA was isolated from leaves using a urea gDNA extraction method (*6*). Genomic DNA was digested by *Bst*1 (NEB) overnight and cleaned with a MinElute DNA clean-up kit (Qiagen). End repair was carried out by incubating overnight with T4 DNA polymerase (NEB) followed by A-tailing before clean-up with a MinElute DNA cleanup kit (Qiagen). To generate the Splinkerette adapter, Long-strand adaptor and Short-strand adaptor oligos were synthesised and annealed by heating for 10 mins at 72ºC and allow them to cool at room temperature. Fragmented DNA was ligated to a Splinkerette adapter by T4 ligase (NEB) followed by clean-up with MinElute DNA cleanup kit (Qiagen). Ligated genomic DNA was then used for two rounds of nested PCR with Phusion High-Fidelity Polymerase (NEB). Splink1 primer and retrotransposon-specific round-1 primer was used for the first round of PCR. This PCR product was used as template for the second PCR with Splink2 primer and retrotransposon-specific round 2 primers. Each PCR product was resolved on a 6% Acrylamide gel and stained with ethidium bromide before imaging.

For the high-throughput sequencing to identify and map retroelement insertion sites we followed the method developed by Dooner, Wang, Huang, Li, He, Xiong and Du (*7*) with minor modifications. An equal amount of young leaf-tissue was harvested from 10 plants and DNA was extracted by the Urea method. A modified Splinkerette-PCR was used to isolate the retroelement insertion sites. DNA was sheared using a Bioruptor sonication system (Diagenode, Belgium) to a mean size of ~1.7-kb and size-selected by 0.8xAgencourt AMPure XP beads (Beckman Coulter, Brea, CA). The protocol of KAPA library preparation Kits (Kapa Biosystems Inc., Wilmington, MA) was followed in subsequent end-repairing, A-tailing and adaptor-ligation procedures. PCR amplifications followed the protocol of Phusion High-Fidelity Polymerase (NEB, Ipswich, MA). Biotin-Splink1 primers were used for 1st round PCR using Phusion High-Fidelity Polymerase (NEB, Ipswich, MA) and the PCR product was purified using Dynabeads® M-280 Streptavidin (Thermo Fisher Scientific, Carlsbad, CA). These purified products were used as template for second PCR using primers for different retroelements that were barcoded to allow sample multiplexing. The oligonucleotides required for Splinkerette-PCR are listed in Supplementary Table S6. We constructed sequencing libraries, after amplicons were end-repaired, A-tailed, and ligated to Illumina TruSeq Single Index Barcoded adaptors from the Illumina TruSeq LT DNA kit (Illumina, Kapa Biosciences). Adaptor-ligated DNA was amplified in a PCR reaction with 1X Kapa HF PCR Master Mix (Kapa Biosciences), and 1X TruSeq PCR Primer Cocktail (Illumina). Libraries were sequenced on an Illumina NextSeq platform (University of Warwick). Primer sequences are listed in Supplementary Table S6. Library details and number of reads are provided in Table S7.

**Sequencing data and statistics analyses**

Analyses of small RNA sequencing data were carried out using previously described methods *(8*). Mapping of small RNAs to AGPv4 reference genome was performed using Bowtie. Any read with more than 50 perfect matches (“hits”) to the genome was excluded from further analysis.

Abundances of small RNAs in each library were normalized to "TP10M" (transcripts per 10 million) based on the total count of genome-matched reads in that library. For the analysis of RNA-seq data, reads were trimmed and mapped to AGPv4 reference genome using TopHat2 (*9*) and the expression annotated genes and transposons was quantified using TEtranscripts (*10*). For the analysis of NanoPARE sequencing data, we used a described analysis pipeline and determined candidates for sRNA-mediated cleavage of retrotransposons using EndCut (*11*).

**Phosphoproteomic analysis**

Protein from anthers were extracted by adding three times the volume of extraction buffer (50 mM HEPES, 150 mM NaCl, 1 mM EDTA, 20 mM NaF, 1 mM Na2MoO4, 1% (v/v) SDS, 1 mM PMSF, 2 µM Calyculin A, 1 mM NaVO4, 1 mM DTT, Protease inhibitor cocktail (Roche)) to 0.5 g of tissue. After 30 min the samples were spun for 15 min at 4,000 g (4°C) to remove debris. The supernatant was transferred to a new tube, centrifuged for 30 min at 16,000 g (4°C), transferred to a new tube and treated using the FASP protocol (*12*). Samples were loaded on Amicon® Ultra-2 mL Centrifugal Filters with a cutoff of 3 kDa and diluted with 1 ml 8 M urea until 1 ml of buffer was passed through the column. Reduction and alkylation of the cysteine residues was carried out by adding a combination of 5 mM tris (2-carboxyethyl) phosphine (TCEP) and 10 mM iodoacetamide (IAA) for 30 min at room temperature in the dark, followed by six washes with 25 mM Hepes (pH 7.5). The protein was digested with trypsin (Promega Trypsin Gold, mass spectrometry grade) overnight at 37 °C at an enzyme-to-substrate ratio of 1:100 (w:w). After digestion the peptides were suspended in 80% acetonitrile (AcN), 5% trifluoroacetic acid (TFA) and the insoluble matter was spun down at 4000 g for 10 min. The supernatant was used for the enrichment of phosphopeptides as previously described with minor modifications (*13*). The peptide concentration was measured with a Qubit™ fluorometer (Invitrogen) and 1 µg total peptides were used for each sample. The Titansphere TiO2 10 µm beads (GL Sciences Inc.) were equilibrated in a buffer containing 20 mg/mL 2,5-dihydroxybenzoic acid (DHB), 80% ACN and 5% TFA in a ratio of 10 µl DHBeq per 1 mg beads for 10 min with gentle shaking at 600 rpm. TiO2 beads were used in a ratio of 1:2 peptide-bead ration (w:w). The TiO2 solution was added to each sample and incubated for 60 min at room temperature. This step was repeated one more time. The samples were then spun down at 3000 g for 2 min and resuspended in 100 µL Wash buffer I (10% AcN, 5% TFA). The resuspended beads were added to self-made C8-columns. C8-colums were made of 200 µL pipette tips with 2 mm Empore™Octyl C8 (Supelco) discs. The columns were spun down at 2600 g for 2 min, washed with 100 µL Wash buffer II (40% AcN, 5% TFA) and 100 µL Wash buffer III (40% AcN, 5% TFA). The peptides were eluted from the TiO2 beads with 20 µL 5% ammonium hydroxide and subsequently with 20 µL 20% ammonium hydroxide in 25% AcN. Both eluates were pooled, the volume was reduced to 5 µL in a centrifugal evaporator (20–30 min) and acidified with 100 µL of buffer A (2% AcN, 1% TFA). Samples were desalted with a self-made C18 column (Empore™Octadecyl C18). C18 were made in the same way as the C8-columns. Before adding the samples, the C18-columns were activated with 50 µL methanol and washed with 50 µL AcN and 50 µL buffer A* (2% AcN, 0.1% TFA). Samples were loaded on the C18-column and spun at 2000 g for 7 min. The columns were washed with 50 µL ethyl acetate and 50 µL buffer A* and then eluted consecutively with 20 µL 40% AcN and 20 µL 60% AcN. Samples were then vacuum-dried and prior to MS analysis resuspended in 50 µL buffer A*.

**Mass spectrometry**

Reversed phase chromatography was used to separate tryptic peptides prior to mass spectrometric analysis. Two columns were utilised, an Acclaim PepMap µ-precolumn cartridge 300 µm i.d. x 5 mm 5 µm 100 Å and an Acclaim PepMap RSLC 75 µm x 25 cm 2 µm 100 Å (Thermo Scientific). The columns were installed on an Ultimate 3000 RSLCnano system (Dionex). Mobile phase buffer A was composed of 0.1% formic acid in water and mobile phase B 0.1 % formic acid in acetonitrile. Samples were loaded onto the µ-precolumn equilibrated in 2% aqueous acetonitrile containing 0.1% trifluoroacetic acid for 8 min at 10 µL min-1 after which peptides were eluted onto the analytical column at 300 nL min-1 by increasing the mobile phase B concentration from 3% B to 35% over 40 min and then to 90% B over 4 min, followed by a 15 min re-equilibration at 3% B.

Eluting peptides were converted to gas-phase ions by means of electrospray ionization and analysed on a Thermo Orbitrap Fusion (Q-OT-qIT, Thermo Scientific). Survey scans of peptide precursors from 350 to 1500 m/z were performed at 120K resolution (at 200 m/z) with a 4 × 105 ion count target. Tandem MS was performed by isolation at 1.6 Th using the quadrupole, HCD fragmentation with normalized collision energy of 35, and rapid scan MS analysis in the ion trap. The MS2 ion count target was set to 1x104 and the max injection time was 200 ms. Precursors with charge state 2–7 were selected and sampled for MS2. The dynamic exclusion duration was set to 45 s with a 10ppm tolerance around the selected precursor and its isotopes. Monoisotopic precursor selection was turned on. The instrument was run in top speed mode with 2 s cycles.

**Mass spectrometry data analysis**

A label-free peptide relative quantification analysis was performed in Progenesis QI for Proteomics (Nonlinear Dynamics, Durham). To identify peptides, peak lists were created by using Progenesis QI. The raw data was searched against maize B73 RefGen_4 Working Gene set. Peptides were generated from a tryptic digestion with up to two missed cleavages, carbamidomethylation of cysteines as fixed modifications, oxidation of methionine and phosphorylation of serine, threonine and tyrosine as variable modifications. Precursor mass tolerance was 5 ppm and product ions were searched at 0.8 Da tolerances. Scaffold (TM, version 4.4.5, Proteome Software Inc.) was used to validate MS/MS based peptide and protein identifications. Peptide identifications were accepted if they could be established at greater than 95.0% probability by the Scaffold Local FDR algorithm. Protein identifications were accepted if they could be established at greater than 99.0% probability and contained at least one identified peptide. Proteins that contained similar peptides and could not be differentiated based on MS/MS analysis alone were grouped to satisfy the principles of parsimony. List of differentially accumulated phosphopeptides are listed in Table S4.

**Protein structure modelling**

To model the conserved catalytic domain of MAGO2, we used MODELLER (*14*). First, we scanned the PDB database (http://www.rcsb.org/) to identify proteins with known structure whose sequences could be best aligned with that of MAGO2. This search identified the *K. polysporus* Argonaute (PDB ID 4F1N) (*15*) as a good match for roughly the last two thirds of the MAGO2 sequence. Moreover, three other proteins, namely the human Argonaute1-3, were found to be viable templates covering the rest of the MAGO2 sequence, with varying degree of overlap with KpAGO. Of these, we chose the Argonaute 2, because of the high resolution and crystallographic quality of one of its available structures (PDB ID 4Z4D) (*16*). To use both structure as templates for our modelling, we first aligned them to each other, and then aligned the fitted structures to the sequence of MAGO2. We then used this multiple-template alignment to produce 64 base models of MAGO2, refining the loop regions in each of them twice. Each refinement was repeated 16 independent times, resulting in a total of 1,024 different models. To find the best one, we assessed each model using a high-resolution version of the DOPE (Discrete Optimized Protein Energy) method (*17*), and picked the model with the best score, checking it by hand to ensure it contained no knotted loops or other unphysical structures.

Having produced an initial model, we refined it using molecular dynamics (MD) simulations, to obtain a realistic final structure. All MD simulations were carried out using Amber18 (*18*). To prepare the parameters for the simulations, we first added hydrogens to the pdb file of the model using the pdb4amber and reduce programs (*19*). To ensure the model was properly folded, we decided to run an initial relaxation simulation using implicit solvation, to exploit the speed-up in conformational sampling that this method provides. More specifically, we used the Generalized Born model (*20*) with a set of optimized atomic parameters for proteins (*21*). Thus, to create the topology parameters, we used the ff14SBonlysc force field, which uses the same parameters as the ff99SB force field (*22*) for the backbone, but full quantum-mechanics ones for the side-chains (*23*), and which is known to work best in this setup.

We then minimized the structure with 16000 steps of steepest descent, before heating the system gradually over 0.5 ns from 0 K to 295.15 K, using a Langevin thermostat with collision frequency of 2.0 ps-1. For this and subsequent implicit-solvation steps we constrained the length of the bonds with hydrogens using SHAKE (*24*), imposed a cutoff for nonbonded pair and effective Born radii calculations of 24 Å, and used an integration step of 2 fs. Also, forces involving the derivatives with respect to the effective Born radii were computed every 2 integration steps. After heating the system, it was allowed to relax at constant temperature, and computed the total potential energy and its individual contributions (bond energy, dihedral angle energy, van der Waals 1-4 interaction energy, electrostatic 1-4 energy, total van der Waals interaction energy and total electrostatic energy). We stopped the relaxation when we observed at least 20 ns of stability in each of the components, as well as in the total potential energy, which we took as an indicator that no further conformational changes were likely to occur (*25*).

To obtain a more realistic model we then performed one more relaxation in explicit solvent. We prepared the starting topology from the last simulation frame of the previous step using the same procedure described above. However, this time we solvated the protein using the TIP3P water model (*26*) in a truncated octahedral box imposing a minimum distance between the edges of the box and the atoms of the protein of 8 Å. Also, for this simulation step we used the full ff14SB force field (*23*), as the current gold-standard for simulations with explicit water molecules, and neutralized the charge of the protein adding 16 Cl- ions, treated via the parameters by Joung and Cheatham (*27, 28*). We then found the optimal distribution of water molecules by constraining the protein atoms via a harmonic potential with a coupling constant of 500 kcal/(mol Å2) and minimizing the potential energy of the system. Subsequently, we removed the constrains and minimized the whole system again, allowing every atom to move. For both minimization steps we used steepest descent and stopped the minimization process when the root-mean-square of the components of the potential energy gradient became smaller than 0.05 kcal/(mol Å2). For this and all other explicit-solvation steps, we used a nonbonded interaction cutoff of 8 Å, constrained the hydrogen-involving bonds using SHAKE, used an integration step of 2 fs, and evaluated slowly-varying terms in the force field at every step.

Having obtained a minimized structure, we heated it using the same protocol described above. Then, we equilibrated the system at a constant temperature and constant pressure of 1 bar for 4.5 ns using a Monte Carlo barostat with pressure relaxation time of 1.0 ps and attempting a volume-change move every 100 integration steps. Finally, we relaxed the system with the same protocol as above, before annealing it to 0 K by decreasing the temperature gradually over 5 ns and minimizing the resulting structure.

To produce a model for the phosphorylated MAGO2 protein, we started from the final non-phosphorylated structure we obtained. We then mutated the relevant residues to their phosphorylated versions, and produced a topology for MD simulations. Since we started this step from an already realistic model of the protein, we had no need to perform an implicit-solvation step before passing to an explicit-solvent simulation. Thus, the protocol we used is the same as the one described above for the explicit-solvation case, with the key changes that we used the phosaa10 force field for the phosphorylated residues (*29, 30*), and the ff99SB for the rest of the protein. This last choice was due to the fact that phosaa10 uses the same assumptions as ff99SB. Thus, its use would not be compatible with force fields of the ff14SB family. Also note that due to the extra negative charges of the phosphoryl groups, only 8 chlorine counter-ions were needed to neutralize the total charge of the system. We then produced a relaxed structure of the system using the exact same steps described previously. To compute the electrostatic potential surfaces, we used PBSA to solve a linearized version of the Poisson-Boltzmann equation, using a level-set-function implementation of the dielectric interface, imposing a smooth molecular surface via density function calculation (*31*), and estimating the nonpolar free energy of solvation as the sum of a cavity term and a dispersion term. For this calculation, we considered an ionic strength of 150 mM, a solvent probe radius of 1.4 Å and a solvent-accessible arc resolution of 0.25 Å.

### 2. **Figures:**

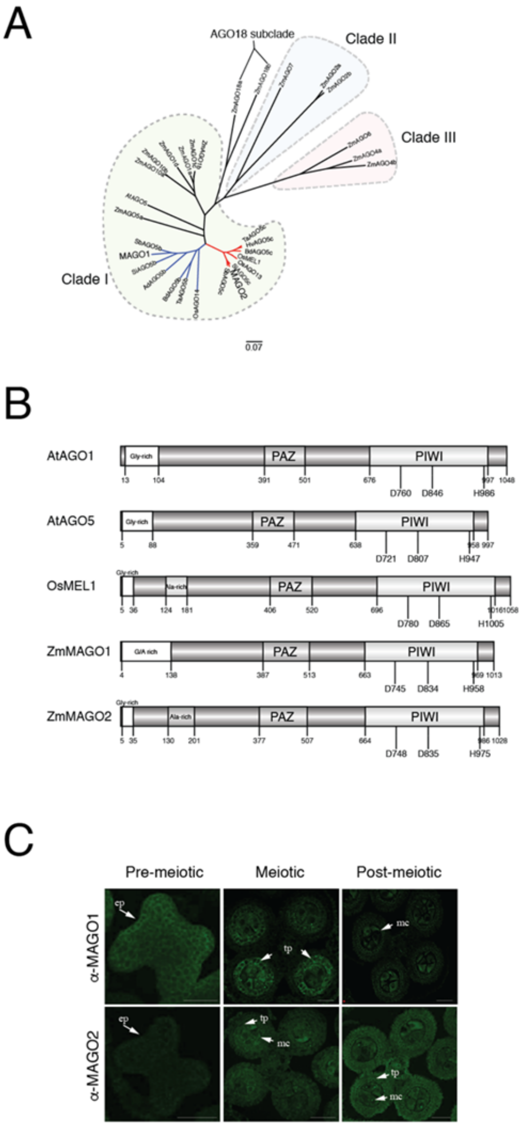


**Fig. S1. Identification of two Male-Associated Argonaute-like (MAGOs) in maize.**
(A) Distance-based phylogeny tree between MAGOs and other monocotyledonous Argonaute-like proteins constructed using the Neighbour-joining method.

(B) Schematic diagram showing the conserved domains present in MAGO and related Argonaute-like proteins. PAZ, Piwi Argonaute and Zwille domain; PIWI, PIWI domain. Catalytic amino acid residues (DDH) are indicated.

(C) Immunodetection of MAGO1 and MAGO2 in developing anthers using specific antisera. White arrow, accumulation of MAGO protein; ep, epidermis; tp, tapetum; mc, meiocyte. Scale bars are 50 μm.

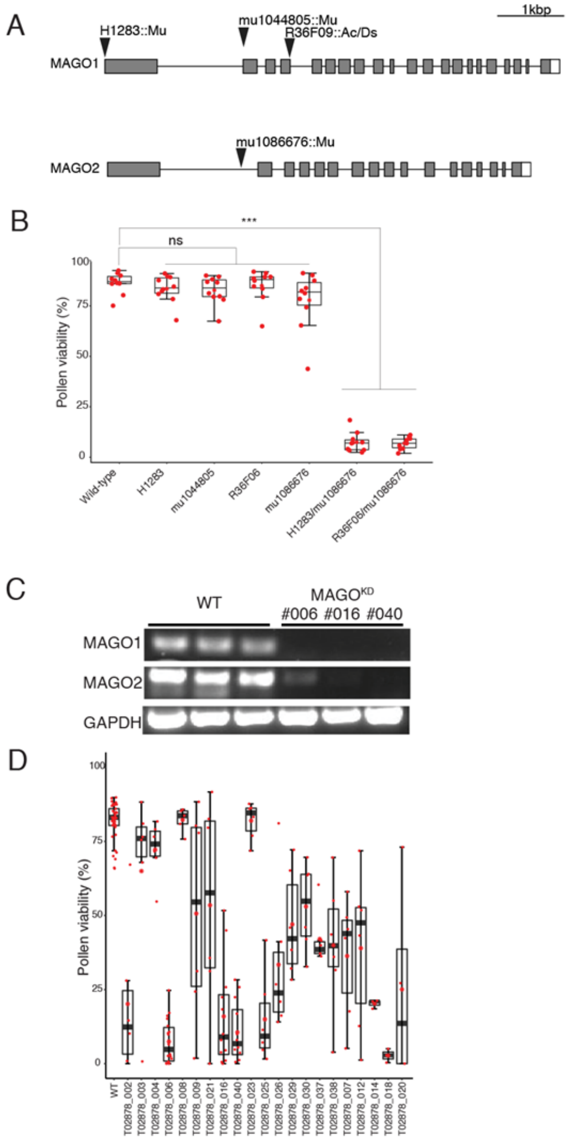


**Fig. S2. Identification and characterization of transposon insertions for MAGO1 and MAGO2 and down-regulation RNAi lines (MAGO$^{KD}$).**

(A) Schematic diagram showing four independent transposon insertion mutant alleles identified for MAGO1 and MAGO2. Black arrowhead, transposon insertion; Grey box, exon; White box, untranslated region.

(B) Pollen viability in wild-type and homozygous transposon insertion plants grown under field conditions. n≥ 10 independent plants analysed per genotype. Differences between groups were determined by Tukey HSD, ***$p < 0.001$; n.s. no-significant.

(C) Accumulation of MAGO 1 and 2 transcripts in pre-meiotic anthers determined by RT-PCR. GAPDH was used as a constitutive control.

(D) Pollen viability in field-grown wild-type and 21 independent MAGO$^{KD}$ lines. n≥ 6 hemizygous T2 plants analysed per genotype; more than 10 anthers analysed per plant. Black line, median; Red star, mean.

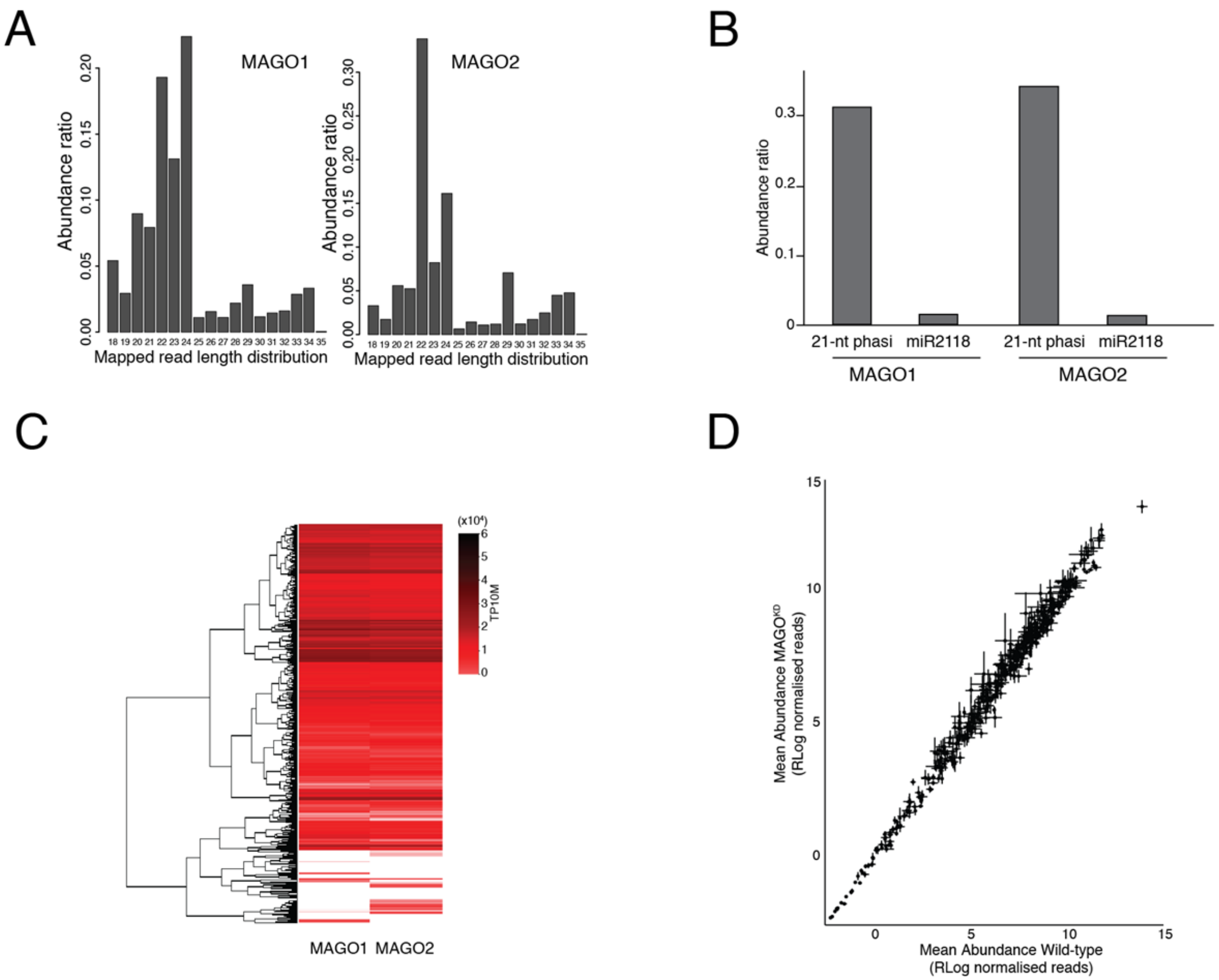


**Fig. S3. Identification of small RNAs associated with MAGO1 and 2.**

(A) Abundance of different sRNA classes in immunoprecipitated MAGO1 and 2 protein fractions determined by sRNA sequencing. n= 2 independent biological replicates. TP10M, transcript per ten million reads.

(B) Abundance of 21-nt phasiRNA and miR2118 trigger in MAGO1 and MAGO2 immunoprecipitated fractions.

(C) Abundance of different 21-nt phasiRNA classes bound to MAGO1 and 2.

(D) Mean abundance plot of 21-nt phasiRNA in pre-meiotic anthers from wild-type and MAGO$^{KD}$ plants. Mean from 3 independent biological replicates; Black line, Standard Deviation.

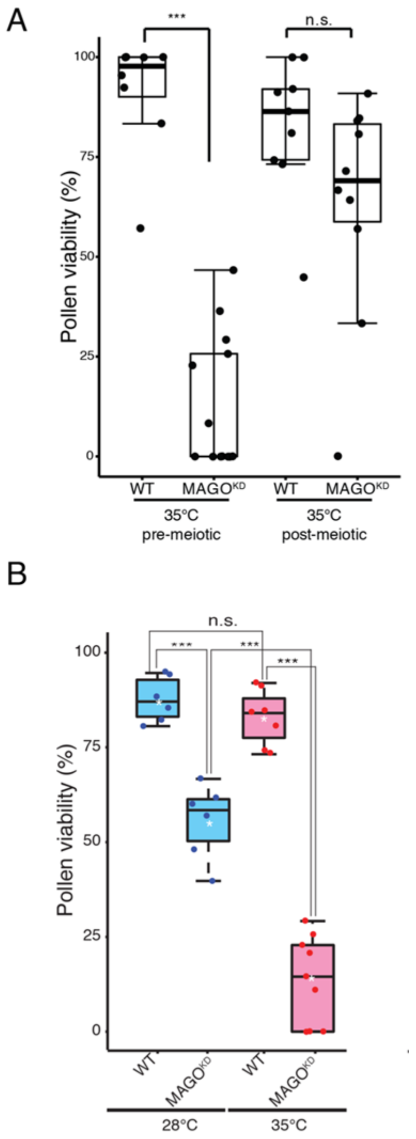


**Fig. S4. MAGO1 and 2 are required before meiosis to sustain male fertility under heat stress.** (A) Pollen viability in wild-type and MAGO$^{KD}$ plants grown under normal conditions (28ºC) and subjected to heat stress (72h/35ºC) before or after meiosis. n≥ 7 plants, more than 6 anthers per plant analysed. Differences between groups were determined by one-way ANOVA, ***p < 0.001; n.s. no-significant.

(B) Pollen viability in wild-type and MAGO$^{KD}$ plants under normal conditions (28ºC) and subjected to heat stress (35ºC) before meiosis (n= 10 plants; 10 anthers each). Differences between groups were determined by one-way ANOVA, ***p < 0.001; n.s. no-significant. Black line, median; White star, mean.

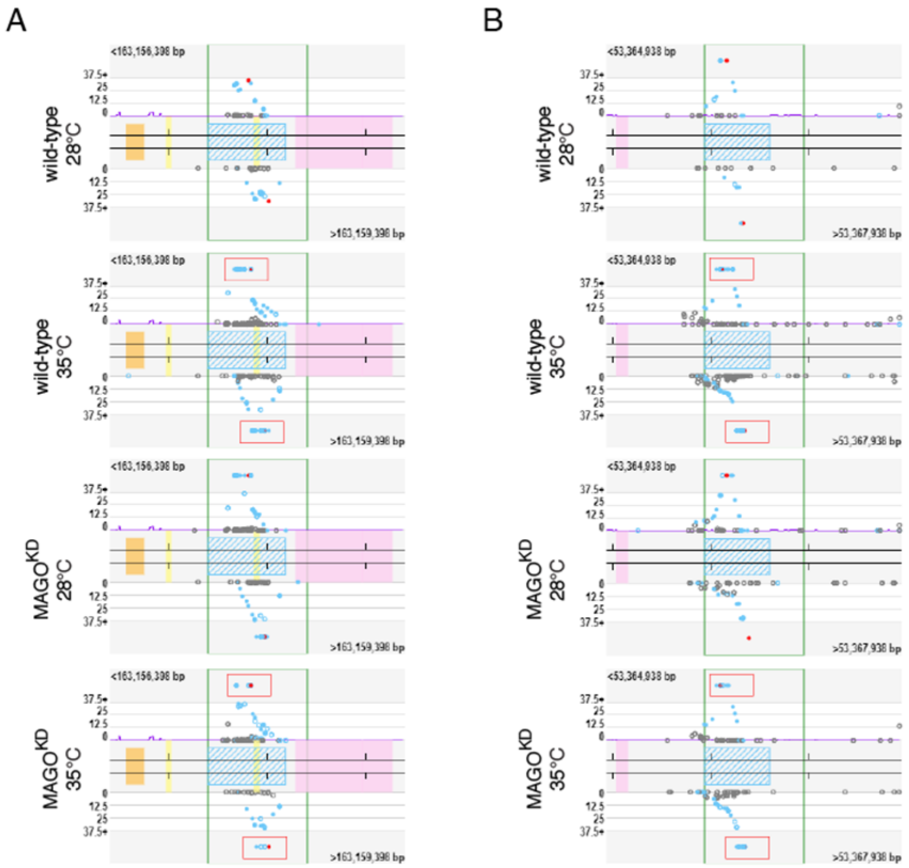


**Fig. S5. Abundance of heat-induced phasiRNAs (Hphasi) from four PHAS loci in pre-meiotic anthers from wild-type and MAGO$^{KD}$ plants grown under normal conditions (28°C) and subjected to heat stress (72h/35°C) before or after meiosis.**

(A) Abundance of 21-nt sRNAs in Hphasi_22 locus. (B) Abundance of 21-nt sRNAs in Hphasi_123 locus. n = 3 independent biological replicates. Red box, Hphasi generating region; Blue dots, common phasiRNAs; Red dot, unique phasiRNAs.

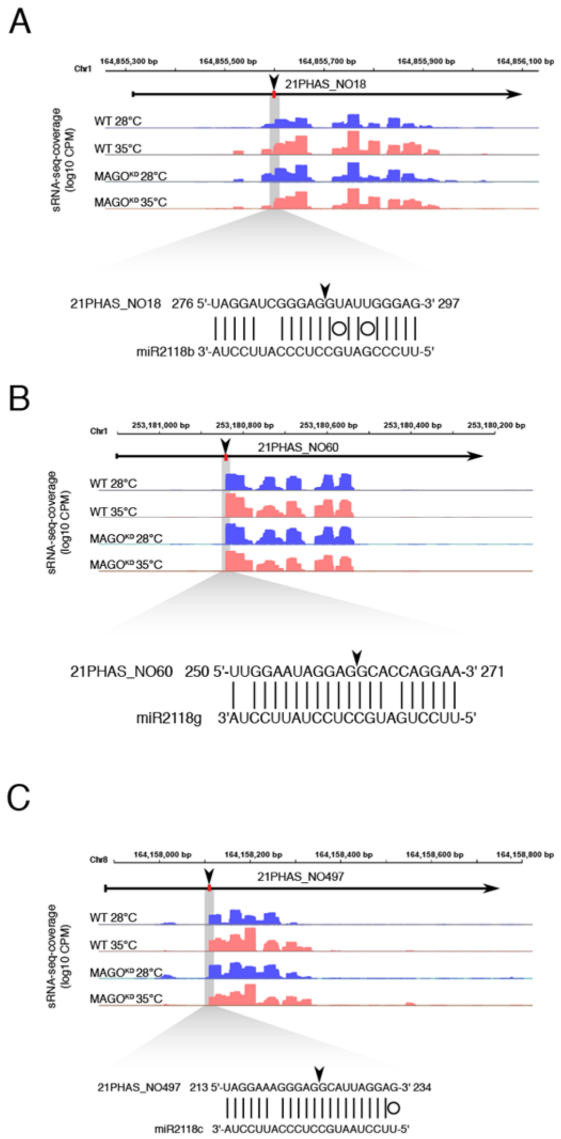


**Fig. S6. Small-RNA-seq coverage of three 21-nt Hphasi generated from miR2118-slicing of different PHAS precursors in wild-type and MAGO$^{KD}$ plants grown under normal conditions (28ºC) and subjected to heat stress (72h/35ºC) before or after meiosis**. n ≥ 3 independent biological replicates. Black arrowhead and red box indicate the location of predicted sites for miRNA-directed slicing remnants.

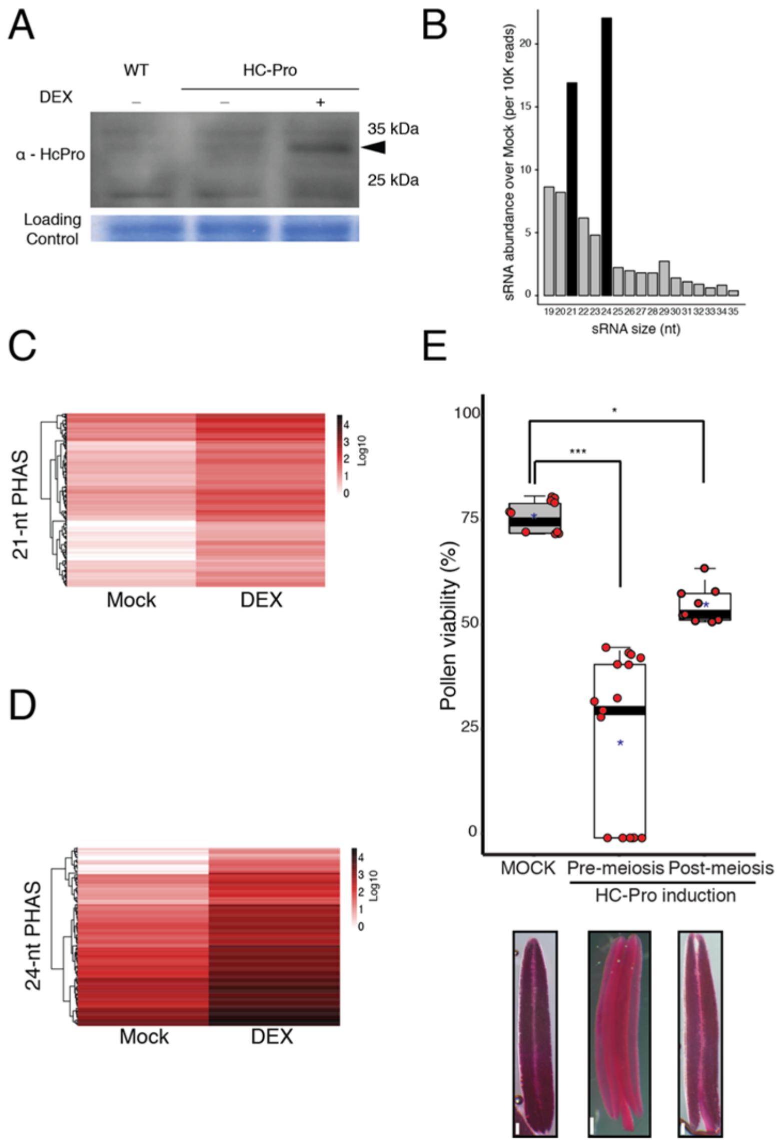


**Fig. S7. Controlled expression of HC-Pro enables the sequestration of small RNAs in maize anthers.**

(A) Western blot detection of HC-Pro accumulation in anthers of wild-type and HC-Pro plants treated with mock and 20 uM DEX.

(B) Relative abundance of sRNAs bound to HC-Pro in pre-meiotic anthers determined by immunoprecipitations coupled to sRNA sequencing. n= 2 independent biological replicates.

(C) Relative abundance of 21-nt phasiRNAs bound to HC-Pro in anthers and determined by immunoprecipitations coupled to sRNA sequencing. n= 2 independent biological replicates.

(D) Relative abundance of 24-nt phasiRNAs bound to HC-Pro in anthers and determined by immunoprecipitations coupled to sRNA sequencing. n= 2 independent biological replicates.

(H) Pollen density in anthers from two independent HDZIV6>>HC-Pro lines (n≥50 anthers; 10 plants each genotype). Differences between groups were determined by one-way ANOVA, ***p < 0.001. Black line, median; Red star, mean. Below, representative anthers after Alexander's Staining. Scale bars 100 μm.

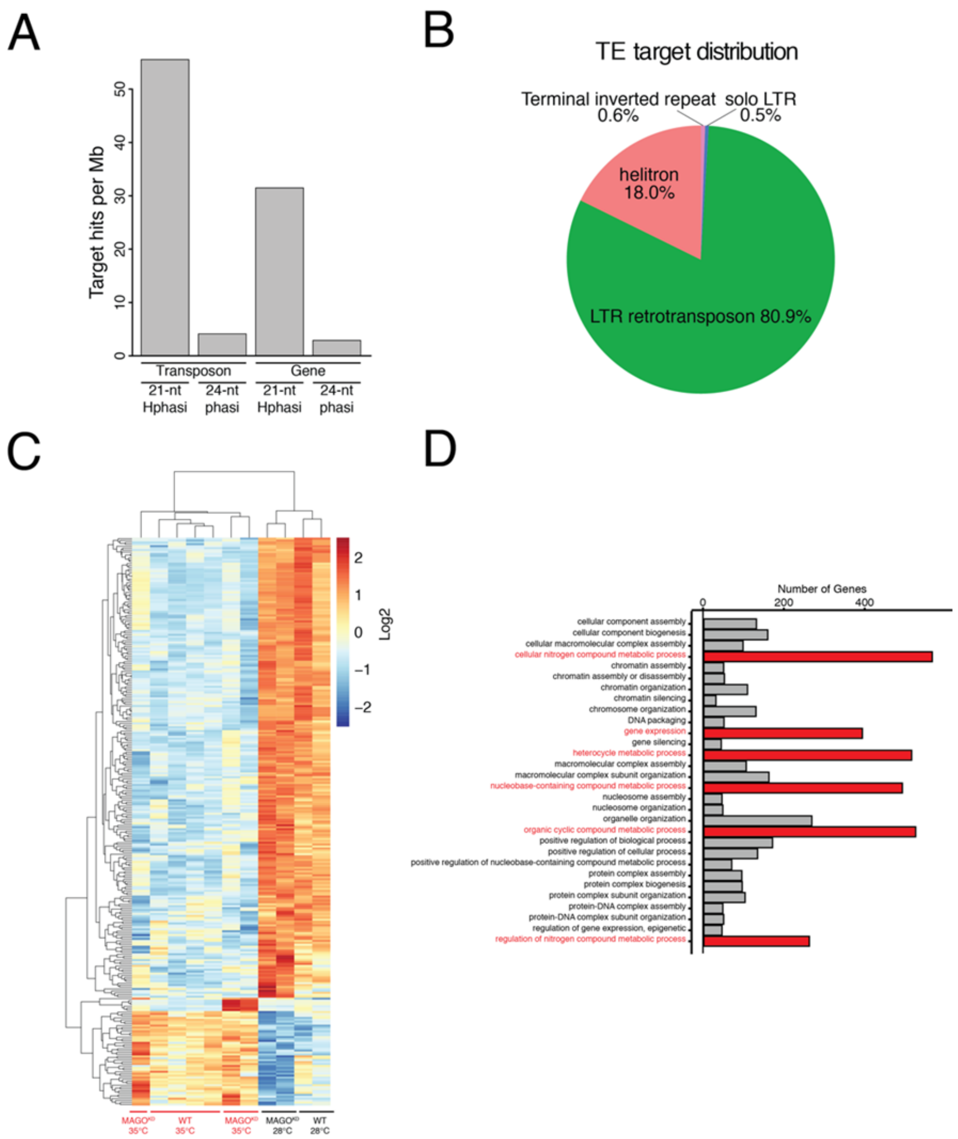


**Fig. S8. Predicted targets of 21-nt Hphasi in the maize genome and impact of heat stress in gene expression.**

(A) Distribution of 21-nt HphasiRNA and 24-nt phasiRNA targets according to their genomic location.

(B) Distribution of 21-nt HphasiRNA targets against annotated transposons.

(C) Heatmap showing transcriptional changes in pre-meiotic anthers caused by heat stress (72h/35ºC). n≥ 2 independent biological replicates**.**

(D) Gene Ontology (GO) analysis showing genes sets enriched within the differentially expressed categories.

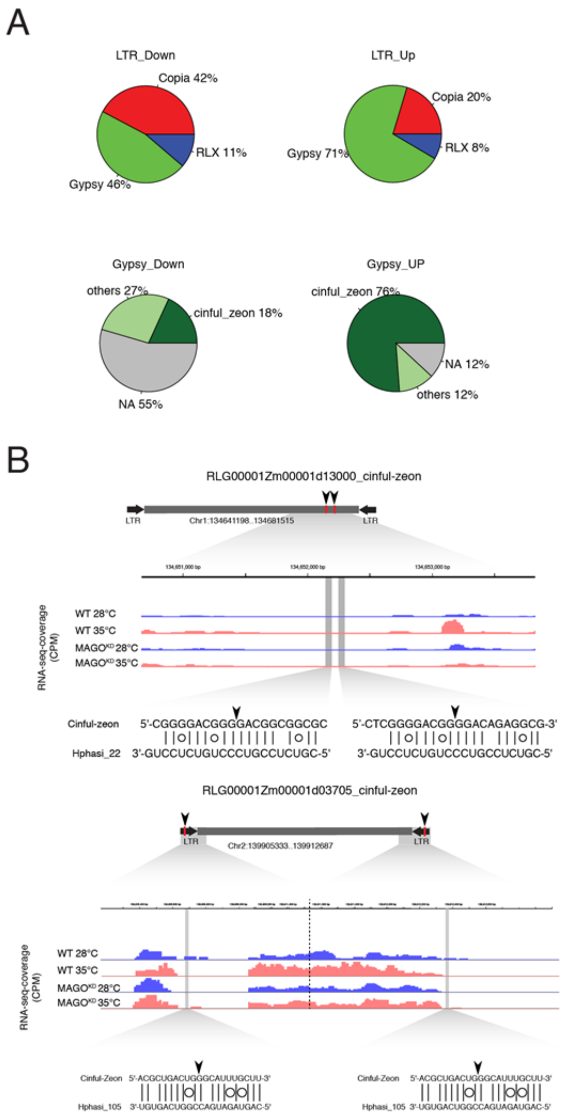


**Fig. S9. Deregulation of retrotransposon in MAGO$^{KD}$ pre-meiotic anthers after exposure to heat stress.**

(A) Frequency of Gypsy, Copia and RLX retrotransposons de-regulated in pre-meiotic anther of MAGO$^{KD}$ plants after exposure to a heat stress (35ºC) and frequency of Cinful-zeon and other retrotransposons of the Gypsy-class de-regulated in pre-meiotic anther of MAGO$^{KD}$ plants after exposure to a heat stress (35ºC).

(B) Coverage of RNA-seq of two different Gypsy-class retrotransposons targeted by Hphasi on anthers of wild-type and MAGO$^{KD}$ plants grown under normal conditions (28ºC) and subjected to heat stress (72h/35ºC) before meiosis. n≥ 3 independent biological replicates**.** Black arrowhead and red box indicate the location of predicted sites for miRNA-directed slicing remnants.

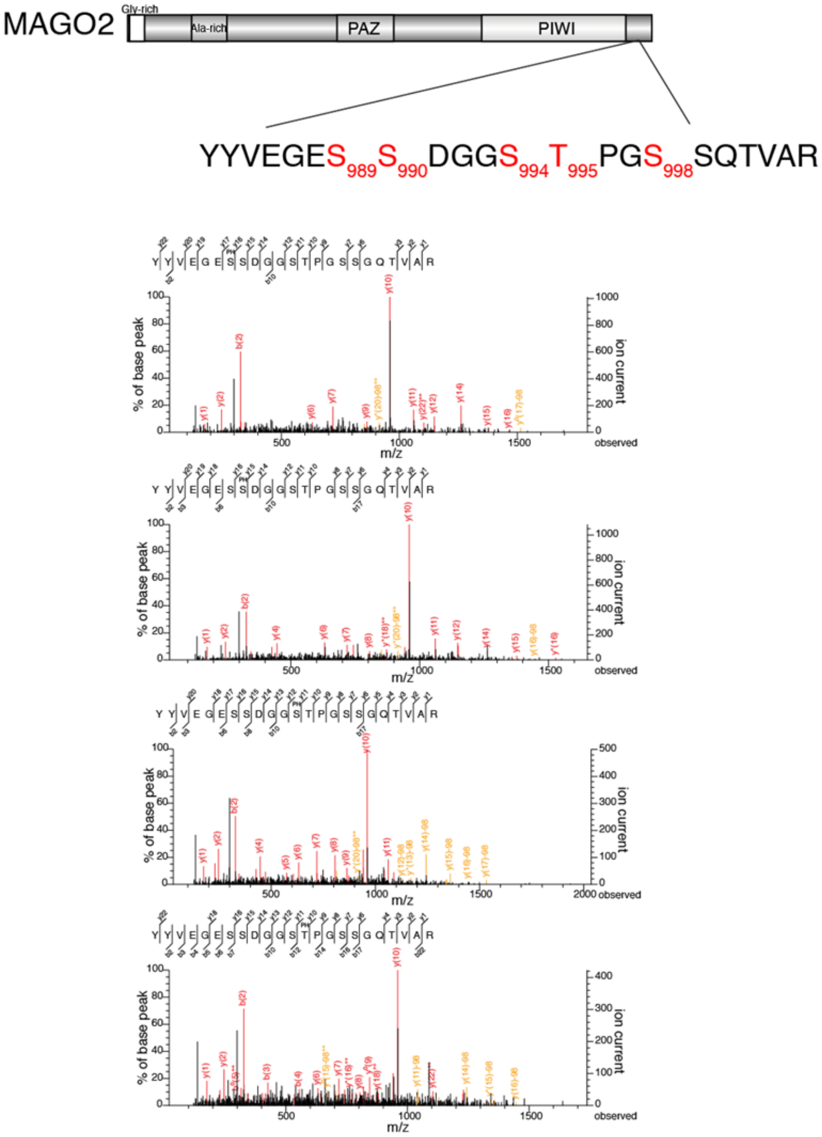


**Fig. S10. MAGO2 phosphopeptides identified by Mass Spectrometry analysis.**
(A) Schematic representation of MAGO2 with annotated serine and threonine residues found to be hypo-phosphorylated in response to heat stress.
(B) Representative spectra of some MAGO2 phosphopeptides.

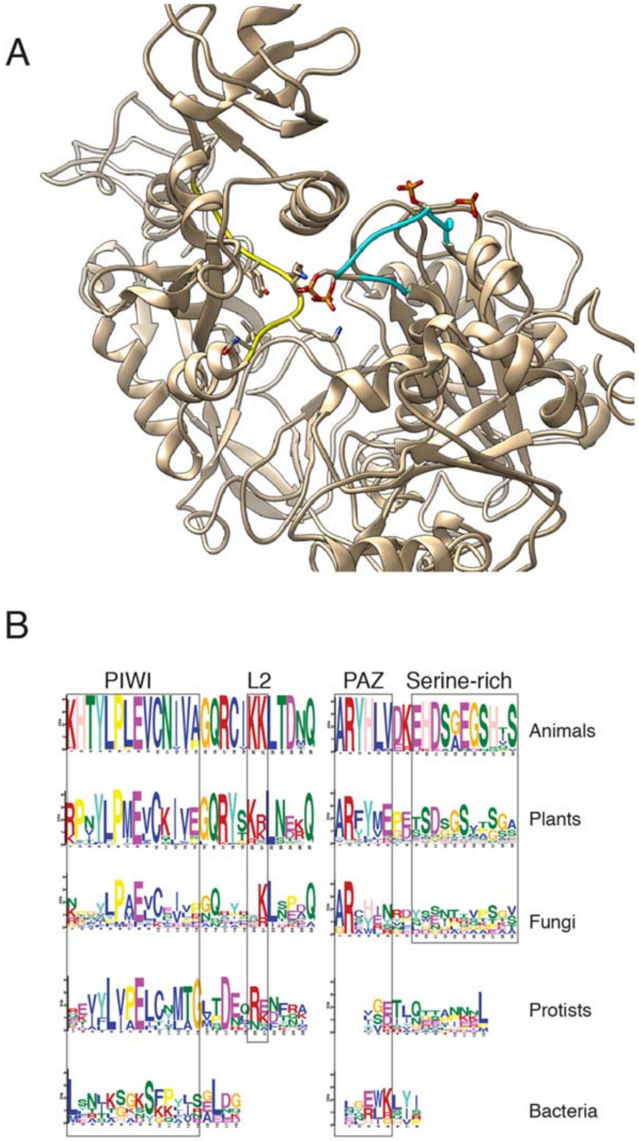


**Fig. S11. Phosphorylation of conserved serine residues in MAGO2 and other catalytically active argonautes.**

(A) Ribbon structure of the PIWI loop and L2 loop of MAGO2 showing the location of dynamically phosphorylated serine (S989, S990, S994 and S998) residues (red) in the PIWI loop (blue).

(B) Amino acid residues conserved in four protein domains of catalytic argonautes.

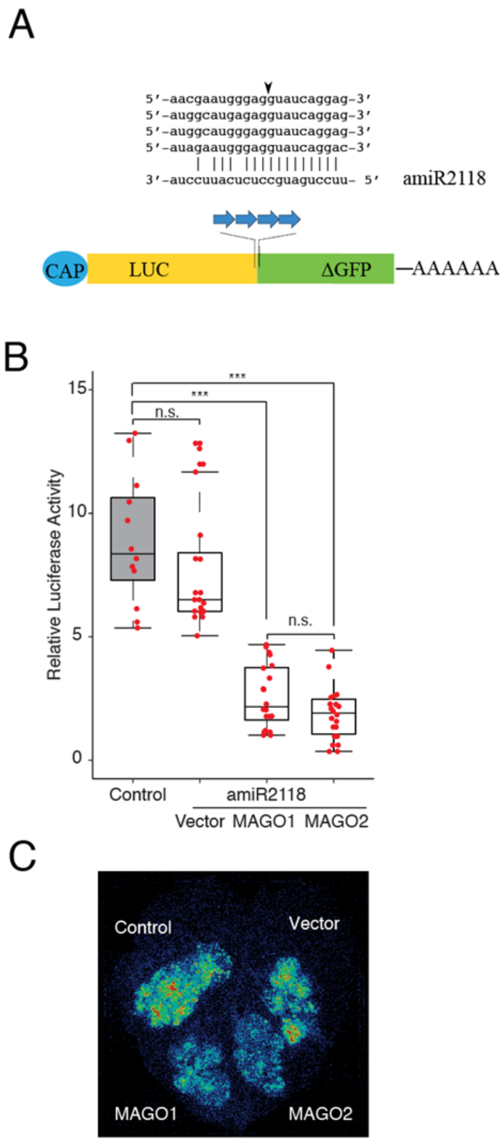


**Fig. S12. *In vivo* silencing activity of MAGO1/2 assessed in *Nicotiana benthamiana* using a Luciferase-amiRNA reporter.**

(A) Schematic diagram of the MAGO sensor construct created by inserting four copies of a miR2118 binding sites in the luciferase (LUC) gene. The predicted cleavage of the target sequences by amiR2118 is indicated by an arrowhead.

(B) Relative luciferase activity in leaves transformed with a control LUC reporter and in combination with amiR2118 and MAGO1 and 2. n= 10 independent biological replicates. Differences between groups were determined by one-way ANOVA, ***$p < 0.001$; n.s. no-significant. Black line, median; Red star, mean.

(C) Bioluminescent imaging of a representative leaf transformed with the LUC reporter used to quantify MAGO1 and 2 silencing activity.

### 3. Tables:

**Table S1.** List of differentially expressed genes in pre-meiotic anthers from wild-type and $MAGO^{KD}$ plants.
**Table S2.** List of predicted LTRs targeted by Hphasi.
**Table S3.** List of new transposon insertions determined by LTR-sequencing.
**Table S4.** List of differentially accumulated phosphopeptides from pre-meiotic anthers of wild-type plants exposed to heat stress.
**Table S5.** Conservation of phosphorylated serine and threonine residues in different Argonaute-like proteins.
**Table S6.** List of oligonucleotides and synthetic DNA constructs.
**Table S7.** Next-Generation-Sequencing library details.

### 4. Multimedia Files:

**Supplementary Movie. Electrostatic potential distribution on the molecular surface of the central cleft of native and phosphorylated MAGO2.** Charge: negative (red), positive (blue) and hydrophobic (grey) residues.
